# Identification and biochemical characterization of a novel eukaryotic-like Ser/Thr kinase in *E. coli*

**DOI:** 10.1101/819920

**Authors:** Krithika Rajagopalan, Jonathan Dworkin

## Abstract

In bacteria, signaling phosphorylation is thought to occur primarily on His and Asp residues. However, phosphoproteomic surveys in phylogenetically diverse bacteria over the past decade have identified numerous proteins that are phosphorylated on Ser and/or Thr residues. Consistently, genes encoding Ser/Thr kinases are present in many bacterial genomes such as *E. coli*, which encodes at least three Ser/Thr kinases. Here we identify a previously uncharacterized ORF, *yegI*, and demonstrate that it encodes a novel Ser/Thr kinase. YegI lacks several conserved residues including those important for Mg^2+^ binding seen in other bacterial Ser/Thr kinases, suggesting that the consensus may be too stringent. We further find that YegI is a two-pass membrane protein with both N- and C-termini located intracellularly.

## Introduction

Reversible protein phosphorylation is an important regulatory mechanism in eukaryotes and prokaryotes (1). In eukaryotes, signaling phosphorylation typically occurs on serine, threonine or tyrosine residues and is mediated by the combined action of kinases and phosphatases. In prokaryotes, signaling phosphorylation has been thought to occur largely on histidine and aspartate residues mediated by histidine kinases of two-component systems (2). However, mass spectrometry based-phosphoproteomic analyses over past decade have identified numerous Ser/Thr/Tyr phosphorylated proteins in many bacteria, including *Escherichia coli* (3–6). Some of these phosphoproteins and their specific phosphosites are conserved in divergent species (4) suggesting that this regulation is physiologically relevant.

Ser/Thr kinases from phylogenetically diverse bacteria have been described (7). Bacterial Ser/Thr kinases can be divided into two classes based on their structure and sequence homology: eukaryotic-like Ser/Thr kinases (eSTKs) and atypical Ser/Thr kinases. eSTKs bear striking resemblance in structure and function to Hanks family Ser/Thr kinases from eukaryotes (7). Unlike eSTKs, atypical Ser/Thr kinases lack any sequence homology to eukaryotic kinases. So far, three genes (*hipA, yeaG, yihE*) encoding Ser/Thr kinases have been reported in *E.coli* of which HipA and YihE are atypical Ser/Thr kinases while YeaG is an eSTK. YeaG plays a role in nitrogen starvation (8), YihE is involved in the Cpx stress response (9) and cell death pathways (10) and HipA regulates bacterial persister formation by phosphorylating a tRNA synthetase (11,12). However, the authentic *in vivo* substrates of these kinases and/or their proximal activating stimuli are largely uncharacterized, complicating efforts to understand their precise physiological role.

Here, we report that the previously uncharacterized *E. coli yegI* gene encodes a Ser/Thr kinase. We describe the biochemical characterization of YegI, which we demonstrate is an integral membrane sSTK. Despite differences in conserved residues observed in other eSTKs, we show that YegI is an active kinase and is sensitive to the kinase inhibitor staurosporine.Additionally, we show that YegI kinase activation follows second order kinetics, consistent with the proposed oligomerization-dependent activation of eSTKs.

## Results

### YegI is a novel eukaryotic-like Ser/Thr kinase

Our lab previously identified a novel PP2C-like phosphatase in the *E. coli* genome encoded by the *pphC* (*yegK*) gene (13). Often, phosphatases of this class are found proximal to eukaryotic-like Ser/Thr kinases (eSTKs) (7). Inspection of the genome of the *E. coli* B strain REL606 revealed the presence of a 1941bp ORF *yegI*, encoding a 646 amino acid protein predicted to be a Ser/Thr kinase, immediately downstream of *pphC*. Of note, BLAST analysis of *E. coli* with previously identified Ser/Thr kinases including YeaG and HipA does not identify *yegI*. In *E. coli* REL606 and in several pathogenic strains such as *E. coli* O157:H7, *yegI* has a four nucleotide overlap (ATGA) at the 5’ end with the 3’ end of *pphC*, suggesting that the genes are co-transcribed. Interestingly, in *E. coli* K strains (*e.g*., MG1655, W3110), the ORF *yegJ*, encoding a protein of unknown function, is found between *yegI* and *pphC* (Supp. Fig. 1) suggesting that it may disrupts co-ordinated expression of *pphC* and *yegI*.

Amino acid sequence analysis and alignment of YegI with bacterial eSTKs (Fig. 1) revealed the presence of Hanks signature motifs (I-XI) suggesting that YegI is an authentic eukaryotic-like Ser/Thr kinase (eSTK) (14). However, YegI contains only seven out of the ten absolutely conserved residues commonly found in eSTKs. For example, in motif III, YegI has an aspartic acid residue instead of the conserved glutamic acid (Fig. 1). This conserved glutamic acid residue contacts Lys-39 in Motif II and is required for stable binding of α and β phosphates of ATP (15). YegI also lacks the conserved asparagine (replaced by serine) in motif VI that together with catalytic site residues Asp-141 and Asp-159 is important for stability of Mg^2+^ ion binding (15). Lastly, YegI lacks a conserved arginine (replaced by glycine) in motif XI that is important for stabilizing substrate-protein interactions (Fig. 1).

**Figure 1.**
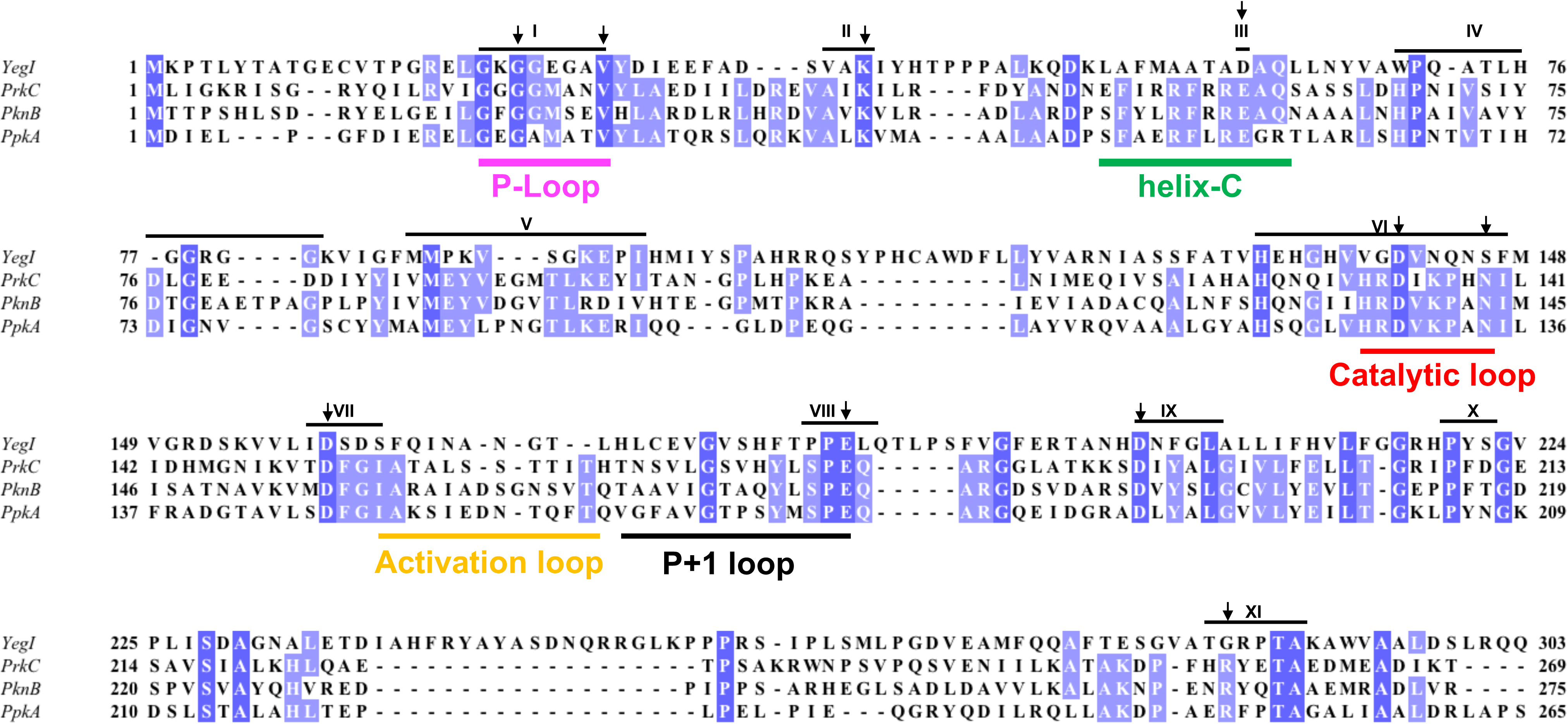
Amino acid sequence alignment of YegI with bacterial eSTKs. YegI (*Escherichia coli*) was aligned to bacterial eSTKs PrkC (*Bacillus subtilis)*, PknB *(Mycobacterium tuberculosis)* and PpkA (*Pseudomonas aeruginosa)* using T-coffee (43). Alignment was visualized using Jalview. Absolutely conserved residues are shaded in dark blue whereas residues conserved in three out of the four kinases are shaded in light blue. The ten absolutely conserved residues found in bacterial eSTKs are depicted with arrows. Hanks conserved domains are depicted on top of the sequence as roman numerals (I-XI). YegI (aa 1-303), PrkC (aa 1-269), PknB (aa 1-275) and PpkA (aa 1-265) which includes the kinase domain was used for alignment. Structurally and functionally important motifs are indicated below the sequence.

A homology model of the YegI kinase domain (aa1-300) based on *M. tuberculosis* eSTK PknB (PDB 1MRU) as a template using Swiss-model (16) demonstrated that despite the absence of these three highly conserved residues, the predicted structure of the YegI kinase domain shows all the structural features seen in *M. tuberculosis* eSTK PknB (Supplementary Fig. 2A). Specifically, YegI adopts the canonical bilobed structure seen in eSTKs. The N-terminal lobe of the YegI kinase domain contains the ATP binding P-loop and the helix C structure (Supplementary Fig. 2B) which is responsible for proper orientation of the active site with the substrate. The C-terminal lobe includes the catalytic loop containing all the residues required for catalysis and an activation loop that is important for kinase activation by autophosphorylation. Thus, although the YegI sequence is only ~20% identical to PknB (Fig. 1), the similarity of the predicted structure of YegI to that of PknB (Supplementary Fig. 2A), strongly suggests that YegI is an eSTK.

### YegI is an active kinase

We previously demonstrated that a His_6_-tagged version of YegI is capable of autophosphorylation (13) despite the absence of three of the ten absolutely conserved residues (Fig. 1). To confirm that YegI belongs to the eSTK family, we examined the consequences of mutations in the conserved lysine at position 39 and a conserved aspartic acid site at position 141, which play a crucial role in ATP binding and catalysis, respectively, in this family (17). We generated YegI mutant proteins carrying either lysine to aspartic acid (K39D) and aspartic acid to asparagine (D141N) changes. The mutant proteins were expressed and purified in the same manner as the wildtype YegI. Kinase activity was demonstrated using an *in vitro* radioactive kinase assay using radiolabeled ***γ***-[^32^P]ATP. Mutation of either Lys-39 or Asp-141 resulted in complete loss of autophosphorylation activity as compared to the wildtype protein indicating that both residues are essential for kinase activity (Fig. 2A).

**Figure 2.**
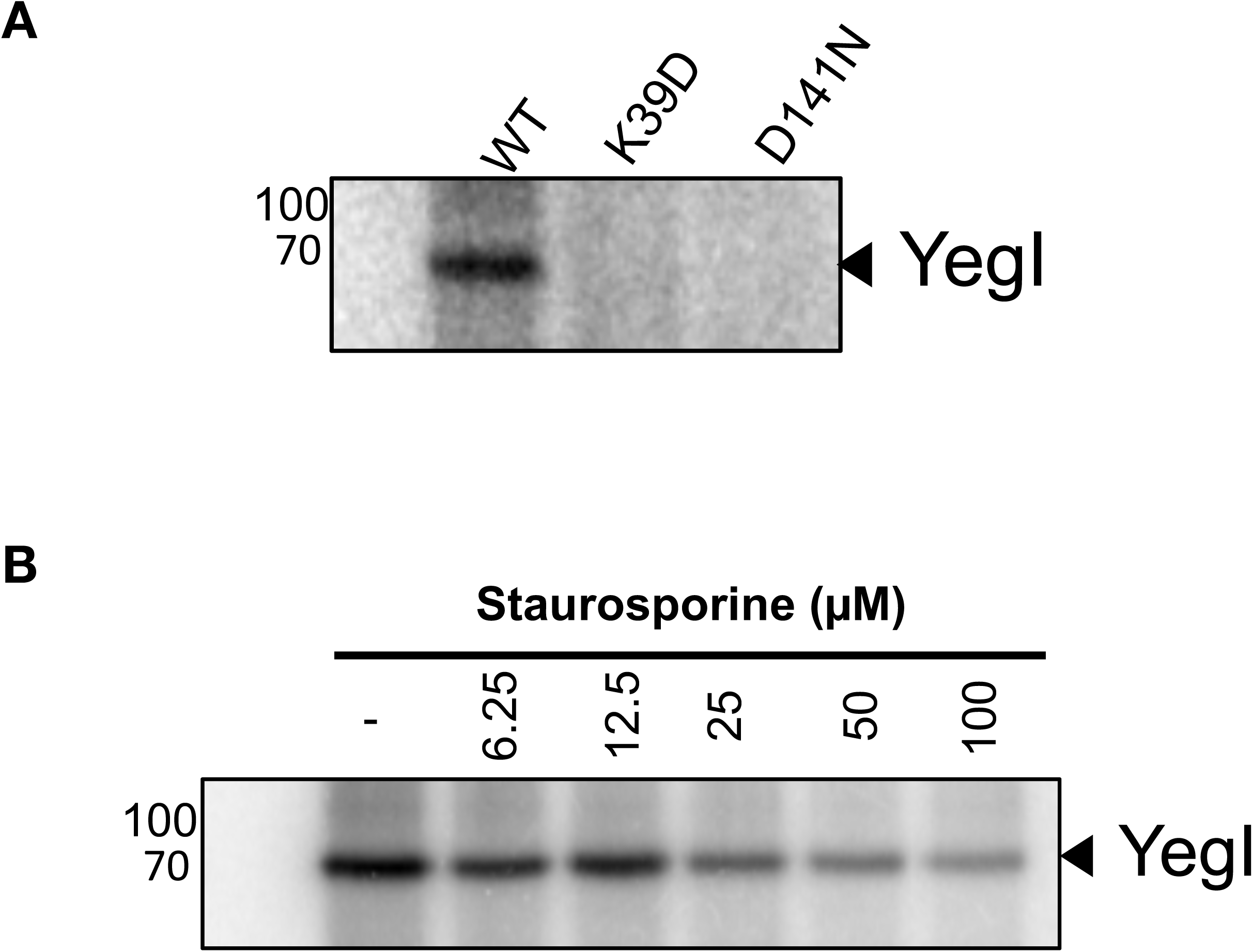
YegI is an active kinase. **(A) Kinase activity of YegI**: Autophosphorylation reactions were carried out at 37 °C with 0.2 μM of YegI (WT or K39D or D141N) in kinase buffer (50 mM Tris pH 7.5, 50 mM KCl, 1 mM DTT, 10 mM MgCl_2_, 10 mM MnCl_2_, 200 μM cold ATP and 5 μCi ***γ***-[^32^P]ATP. Reactions were stopped at t=30 mins and run on 12 % SDS-PAGE followed by autoradiography. Molecular weights (kDa) are indicated on the right of the gel. (B) **Sensitivity to staurosporine:** Autophosphorylation reactions were carried out at 37 °C with 1 μM of YegI (WT) in kinase buffer without cold ATP. Different concentrations of staurosporine (μM) was added to the reaction at indicated concentrations. Reactions were stopped at t=30 mins and run on 12% SDS-PAGE followed by autoradiography. Molecular weights are indicated on the right of the gel.

We examined how the lack of some of the conserved residues affected YegI function in a series of standard assays of bacterial Ser/Thr kinases. For example, staurosporine is widely used as a small molecule inhibitor for Ser/Thr kinases (18) and bacterial eSTKs are typically sensitive to staurosporine with IC_50_ in the μM range (19). Sensitivity of the YegI kinase to staurosporine was assessed by incubating YegI with increasing concentrations of staurosporine. YegI shows an IC_50_ of ~50 μM (Fig. 2B), comparable to the *Enterococcus faecalis* eSTK IreK (IC_50_ of 150 μM (20)) and the *Staphylococcus epidermis* eSTK Stk (IC_50_ of 320 μM (21)). Bacterial eSTKs require the presence of bivalent cations for activity (22–25). We therefore investigated the effect of different bivalent cations on YegI activity by performing radioactive kinase assays in the presence of either MgCl_2_/MnCl_2_/CaCl_2_/NiCl_2_. Similar to many eSTKs, YegI catalyzed autophosphorylation only in the presence of MgCl_2_ and MnCl_2_, albeit to different levels. YegI displayed much higher autophosphorylation in the presence of Mn^2+^ ion than Mg^2+^ ion suggesting that YegI is a Mn^2+^ dependent kinase (Supplementary Fig. 3). The concentration dependence of YegI kinase activity on Mg^2+^ or Mn^2+^ was measured and the optimal MnCl_2_ concentration was determined to be between 0.1-1 mM whereas optimal MgCl_2_ concentration was between 1-10 mM (Fig. 3A) indicating that YegI kinase is more active in the presence of MnCl_2_ than MgCl_2_.

**Figure 3.**
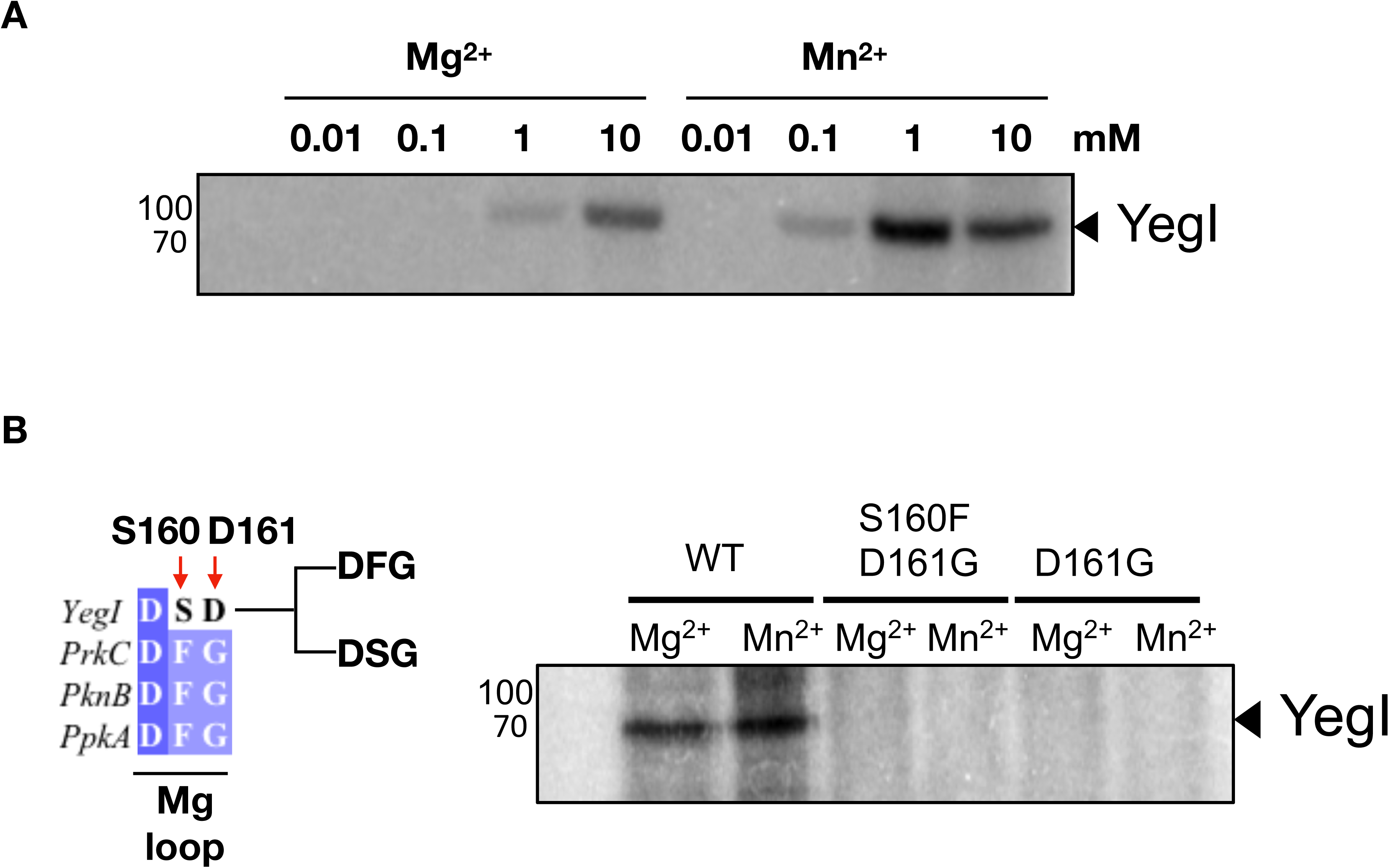
YegI is a Mn^2+^ dependent kinase. **(A) Requirement of Mg^2+^/Mn^2+^ for kinase activity:** Autophosphorylation reactions were carried out at 37 °C with 1 μM of YegI (WT) in kinase buffer with indicated concentrations of MgCl_2_/ MnCl_2_. Reactions were stopped at t=30 mins and run on 12% SDS-PAGE followed by autoradiography. Molecular weights are indicated on the right of the gel. **(B) Requirement of DFG motif**: Autophosphorylation reactions were carried out at 37 °C with 0.2 μM of YegI (WT or S160FD161G or D161G) in kinase buffer with either 10 mM MgCl_2_ or MnCl_2_ as described in Supp. Figure 3

In eSTKs, the highly conserved DFG motif present in domain VII coordinates Mg^2+^ ion binding (26,27). However, amino acid sequence alignment reveals that YegI does not contain the DFG sequence found in bacterial eSTKs but instead the sequence DSD (Fig. 1; Fig. 3B). To examine if addition of the DFG motif to YegI alters preference to MgCl_2_, we generated a mutant protein carrying changes of Ser-160 to a phenylalanine and Asp-161 to a glycine (S160F, D161G). Additionally, we generated a mutant protein carrying a change of Asp-161 to a glycine (D161G) which would change the DSD sequence to the DSG consensus motif. Mutants were expressed and purified in the same manner as wildtype YegI and kinase activity in the presence of MgCl_2_/MnCl_2_ was assessed. Surprisingly, replacement of DSD to DFG or to DSG completely inactivated the kinase (Fig. 3B) indicating that the DSD motif is indispensable for YegI autophosphorylation.

### Autophosphorylation of YegI

Autophosphorylation is a common model for activation of eukaryotic STKs (27), therefore characterization of the kinetics of autophosphorylation and the sites of autophosphorylation can provide insight into how bacterial homologs of these kinases are activated. We performed standard autophosphorylation assay (Fig. 4A) using a range of YegI concentrations and measured formation of radiolabeled phosphorylated YegI (Fig. 4B). Our results indicate that YegI autophosphorylation is not first order and are consistent with YegI undergoing oligomerization. To map the sites of autophosphorylation, His_6_-YegI that had been subjected to an *in vitro* kinase reaction was analyzed by mass spectrometry. Five phosphopeptides that each included at least a single serine residue were identified, leading to the conclusion that Ser-153, Ser-160, and Ser-162 in the predicted activation loop and Ser-602 and Ser-644 at the C-terminus of YegI were phosphorylated (Fig. 5A). To characterize the functional importance of the C-terminal phosphosites, we generated single and double phosphoablative serine to alanine mutants. The mutant proteins were expressed and purified in the same manner as the wildtype protein and assessed for loss of autophosphorylation using a radioactive kinase assay. Since mutation of both Ser-602 and Ser-644 to alanine resulted in only a modest decrease in overall autophosphorylation, phosphorylation of these residues is not necessary for normal YegI activity (Fig. 5B).

**Figure 4.**
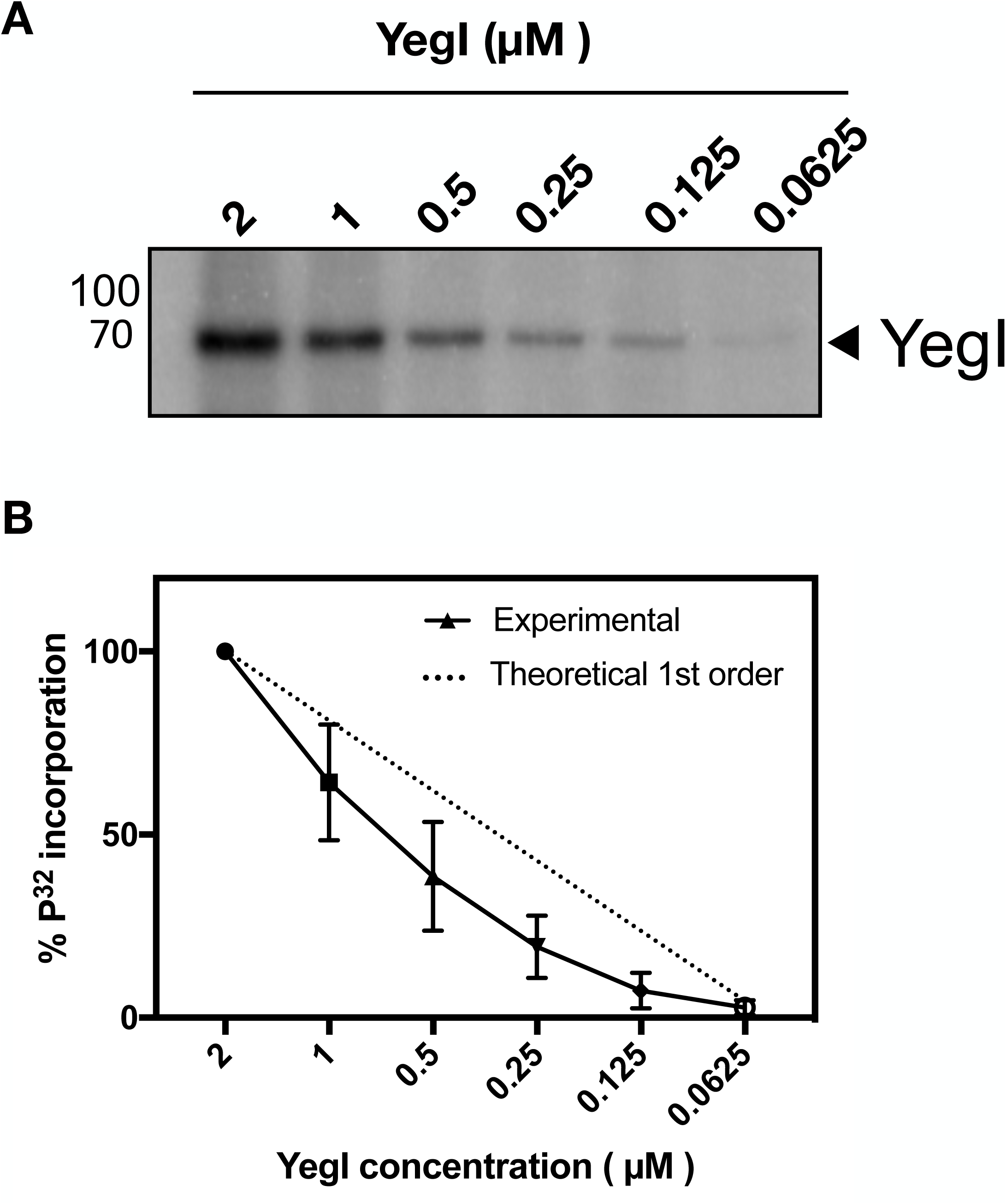
Kinetics of YegI autophosphorylation. (A) **Autophosphorylation of YegI**: Autophosphorylation reactions were carried out at 37 °C with indicated concentrations of YegI (WT) in kinase buffer. Reactions were stopped after 30 min and run on 12% SDS-PAGE followed by autoradiography. Molecular weights are indicated on the right of the gel. (B) **Autophosphorylation of YegI follows second order kinetics**: Kinetics of YegI autophosphorylation was assessed using different concentrations of YegI. % P32 incorporation was calculated relative to autophosphorylation in the presence of 2 μM of YegI from Fig 4(A).

**Figure 5.**
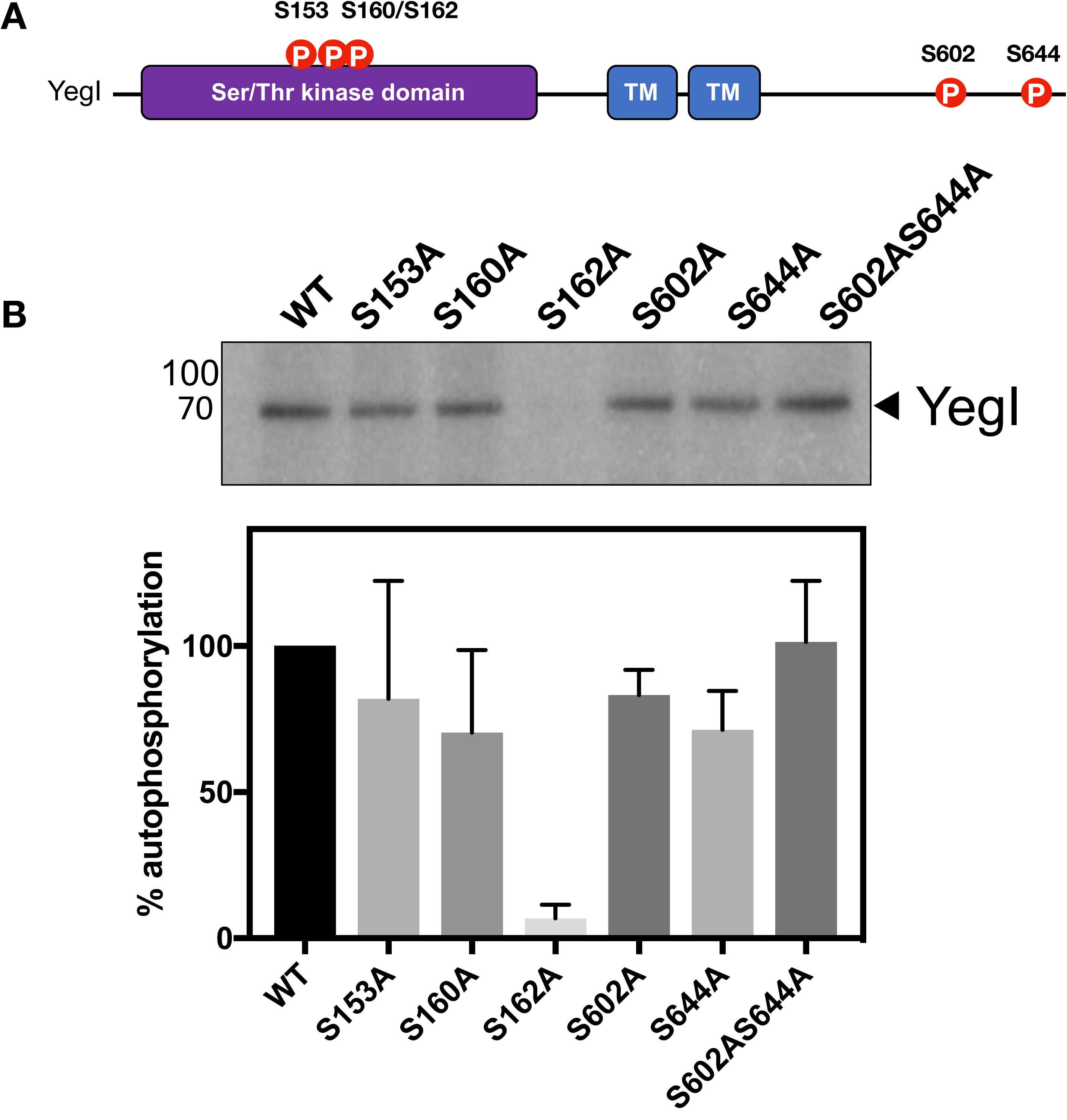
YegI undergoes autophosphorylation on serine residues in the kinase domain and the C-terminus. (A) **Graphical representation of YegI phosphorylation sites:** Domain structure depicting phosphorylation sites following mass spectrometry analysis of autophosphorylation. Phosphorylated residues are indicated by red circles. (B) **Sites of autophosphorylation on YegI**: Reactions were carried out at 37 °C with 1 μM of YegI (WT or phosphoablative mutants) in kinase buffer. Reactions were stopped at t=30 mins and run on 12 % SDS-PAGE followed by autoradiography. Molecular weights (kda) are indicated on the right of the gel.

Since eSTKs undergo autophosphorylation on Ser or Thr residues in the activation loop of the kinase domain (28–31), we investigated the phosphosites (Ser-153, Ser-160, Ser-162) in the putative YegI activation loop. All three residues are located between Motif VI and VII in close proximity to the catalytic loop and activation loop suggesting that phosphorylation of these residues could be important for kinase activity. We mutated each Ser to an alanine to examine if phosphorylation of these Ser residues impacted kinase activity. We observed that mutating Ser-162, but not the other phosphosites (Ser-153 or Ser-160), resulted in a complete loss of autophosphorylation (Fig. 5B), demonstrating that autophosphorylation of Ser-162 is important for YegI activation.

### YegI is an integral membrane protein

We observed that purification of recombinant His6-YegI required the presence of the anionic detergent N-lauryl sarcosine suggesting that YegI is a membrane protein. Consistently, the membrane topology algorithm Phobius (32) predicts that YegI consists of two transmembrane domains joined by a short periplasmic loop with both the N- and the C-terminus located in the cytosol (Fig. 6A). We used the PhoALacZα dual reporter system (33–36) to experimentally validate this predicted topology. In this system, alkaline phosphatase (PhoA, aa 21-471) lacking the periplasmic localization signal is fused in frame with the α-fragment of *E. coli* β-galactosidase (LacZα, aa 5-60), creating PhoALacZα. Localization of a protein of interest is then analyzed by fusing it to PhoALacZα. Since PhoA is active only in the periplasm and LacZ is only active in the cytoplasm, the PhoALacZα fusion displays high phosphatase activity but no β-galactosidase activity when targeted to the periplasm or to the membrane. Conversely, the fusion displays high β-galactosidase activity but no phosphatase activity when it is targeted to the cytoplasm. PhoAlacZα reporter fusions can be assessed using dual indicator plates with PhoA or LacZ specific chromogenic substrates that allow identification of periplasmic or cytoplasmic fusions (35). Specifically, cells expressing PhoAlacZα protein fusions with high levels of phosphatase activity process the X-phos (5-bromo-4-chloro-3-indolyl phosphate) substrate yielding blue colored colonies, while cells expressing PhoAlacZα protein fusions with high levels of β-galactosidase activity process the Red-gal (6-chloro-3-indolyl-β-D-galactoside) substrate yielding red colored colonies.

**Figure 6.**
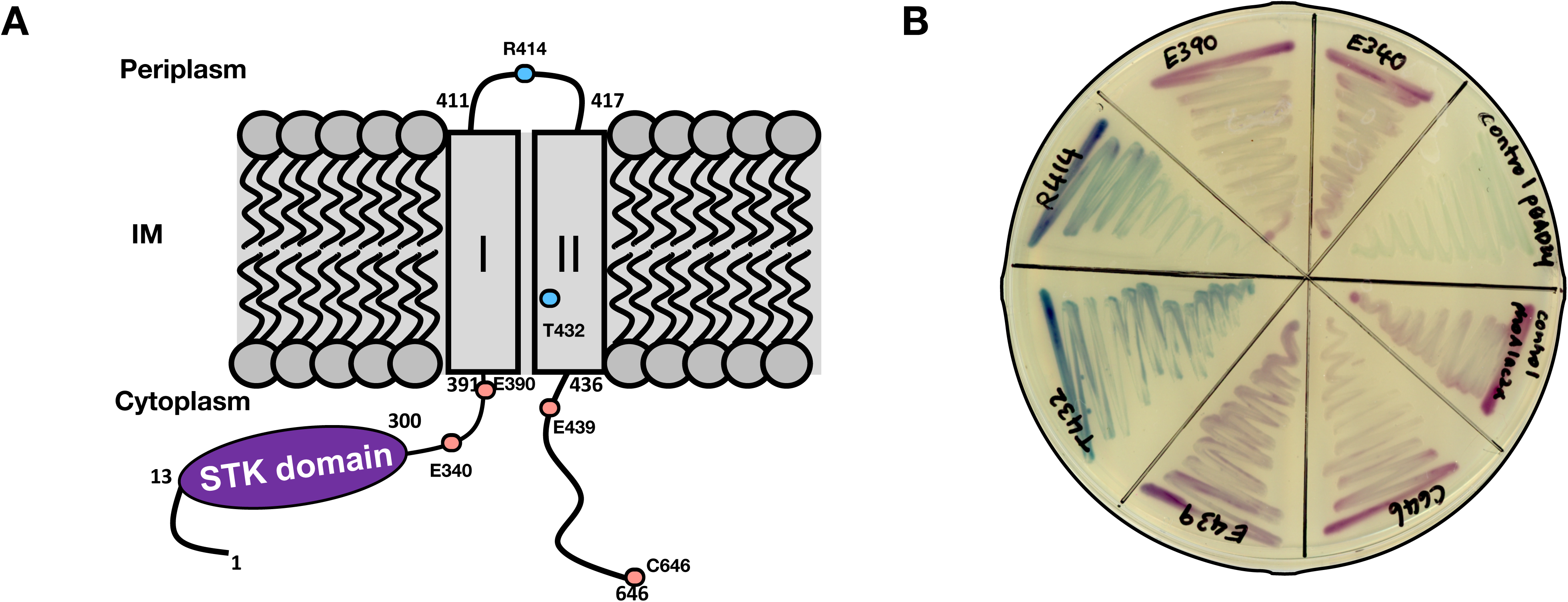
YegI is an integral membrane protein. A) **Predicted topology of YegI using Phobius.** Amino acid positions at which phoA-lacZα reporters are fused are indicated along with associated phenotypes. Phenotypes of phoAlacZα fusions due to LacZ activity are denoted as red ellipses while PhoA activity are denoted as blue ellipses. Predicted transmembrane domains (I and II) are depicted as grey rectangles. (B) **Membrane topology analysis:** DH5α transformed with different phoAlacZα reporters were streaked on an LB plate containing 5-bromo-4-chloro-3-indolyl phosphate disodium salt (X-Phos, 80μg/ml), 6-chloro-3-indolyl-β-D-galactoside (Red-Gal, 100 μg/ml, 0.1 % w/v arabinose and 100 μg/ml of ampicillin. Control pBAD24: control strain expressing pBAD24 empty vector, control phoAlacZα: strain expressing phoAlacZα in pBAD24 without YegI. Amino acid positions at which phoAlacZα reporters are fused are indicated.

We constructed C-terminal PhoALacZα fusions to YegI at amino acid positions 340, 390, 414, 432, 439 and 646 (Fig. 6A) and analyzed them on dual indicator plates containing X-phos and Red-gal. DH5α cells carrying empty backbone vector alone served as a negative control (“vector”) and were colorless, and DH5α cells expressing PhoALacZα without YegI (“PhoALacZα”) served as a positive control and were red/pink indicative of cytoplasmic location (Fig. 6B). Consistent with the *in silico* prediction, PhoALacZα fusions to YegI at amino acid positions 340, 390, 439 and 646 yielded red/pink colored colonies suggestive of high β-galactosidase activity and cytoplasmic localization (Fig. 6B; Table 1). In contrast, PhoALacZα fusions of YegI at amino acid positions 414 and 432 yielded blue colored colonies on indicator plates suggestive of high alkaline phosphatase activity and periplasmic/TM localization (Fig. 6B). In summary, the PhoAlacZα reporter fusions confirmed the bioinformatic prediction that YegI is an integral inner membrane protein containing two TM domains connected by a short periplasmic loop, with both the N- and C-terminus located in the cytosol.

**Table 1.**
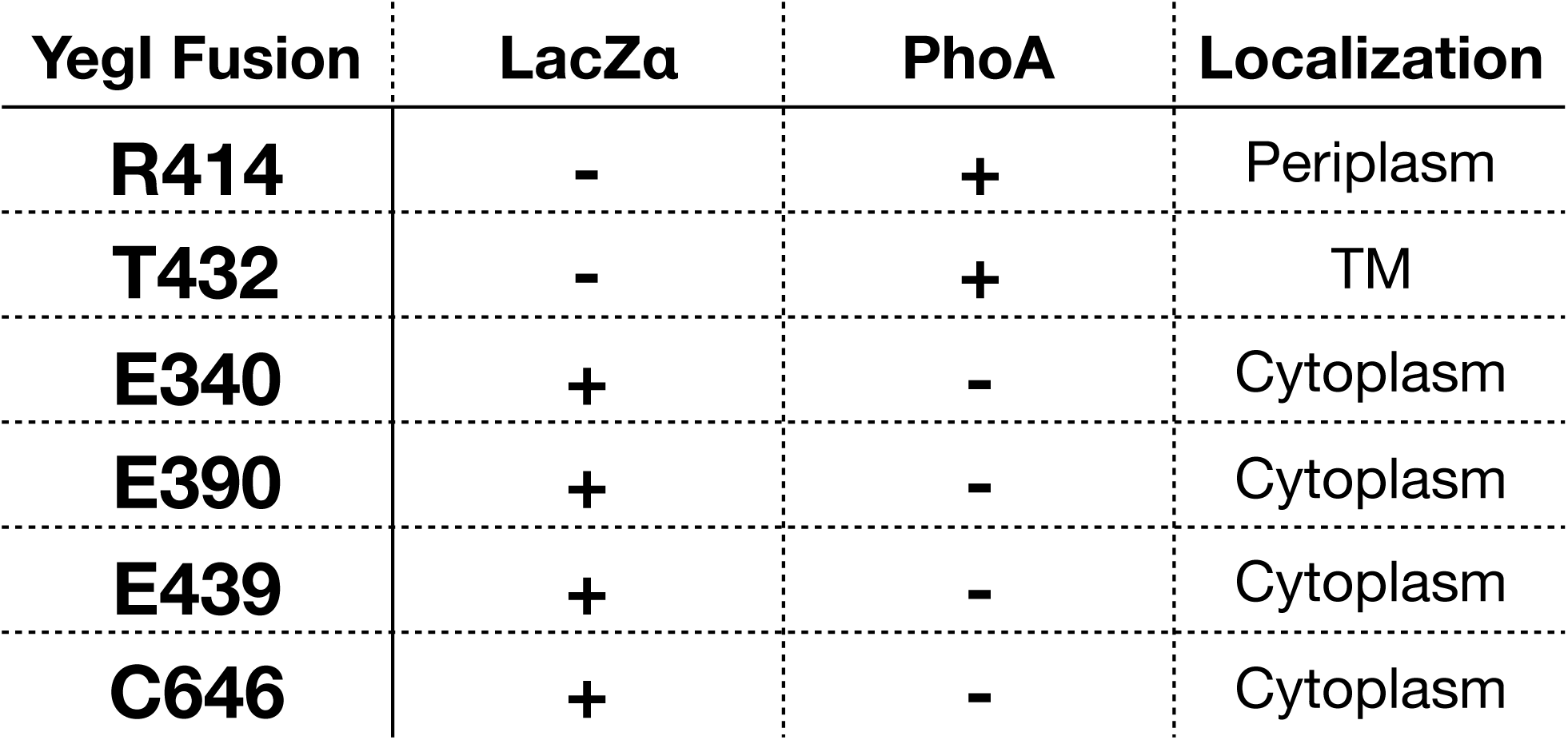
Localization of different phoAlacZα reporter fusions.

## Discussion

Ser/Thr phosphorylation is an important post translational modification that is mediated by Ser/Thr kinases and cognate Ser/Thr protein phosphatases which have been identified in phylogenetically diverse organisms. Although Ser/Thr phosphorylation in the Gram-negative bacterium *E. coli* has been known for >10 years, the enzymes responsible for this reversible modification and their specific substrates are not well understood. In this study, we have described the characterization of YegI, a novel Ser/Thr kinase *E. coli*.

*In silico* analysis revealed that YegI belongs to the class of eSTKs that contain all the 11 key structural subdomains that are characteristic of eukaryotic protein kinases (17). Homology search using YegI kinase domain revealed that it was similar to kinase domains of several bacterial eSTKs (Supplementary Fig. 2A). Unlike other bacterial eSTKs, however YegI includes only seven of the ten absolutely conserved residues responsible for ATP binding, catalysis, and substrate recognition (14,17) (Fig. 1). Despite the absence of these conserved residues, YegI encodes an active protein kinase (Fig. 2A) that undergoes autophosphorylation primarily on serine residues (Fig. 5A) Similar to most bacterial eSTKs, autophosphorylation of YegI on serine residues in the activation loop is critical for catalytic activity of YegI (Fig. 5B).

Although the predicted structure of YegI closely resembled the structure of the bacterial eSTK PknB (Fig. 4.1D), the YegI sequence shows striking deviations in the conserved regions particularly in the catalytic loop and the Mg^2+^-binding DFG loop (Fig. 1). Specifically, the YegI sequence contains the catalytic loop VGDXXQXS as opposed to the canonical catalytic loop HRDXXPXN found in almost all other bacterial eSTKs. The conserved histidine residue in the catalytic loop is crucial for binding both the carbonyl group of catalytic aspartate in the catalytic loop and for making a hydrophobic contact with the phenylalanine in the DFG loop. Additionally, YegI lacks the highly conserved DFG (aspartate-phenylalanine-glycine) loop in motif VII (Fig. 1, 3B) which recognizes the Mg^2+^-bound ATP (37). Instead, YegI contains a DSD (aspartate-serine-aspartate) sequence suggesting either that YegI might be an inactive kinase or that the DFG loop is not essential for every kinase. Our results suggest the latter, since, despite these differences in highly conserved residues, YegI is an active kinase (Fig. 2A) that requires the bivalent cations Mg^2+^/Mn^2+^ for activity with preference for Mn^2+^ (Fig. 3A). We speculated that addition of DFG loop could help kinase activation. However, replacing the DSD loop in YegI with the canonical DFG loop failed to increase activity or preference for Mg^2+^ ion and instead completely abolished activity (Fig. 3B).

This result suggests that the presence of a serine and an aspartic acid residue in the DSD loop might be critical for activity and could affect YegI autophosphorylation. Consistently, we observed that the serine residue in the DSD loop is one of the three serine residues that undergoes autophosphorylation and is required for activity of the kinase (Fig. 5A,B). Substitution of all three serine residues with alanine residues (Ser153Ala, Ser160Ala, Ser162Ala) decreased YegI kinase activity to different extents indicating that these residues are important for autophosphorylation of YegI (Fig. 5B). Particularly, mutation of Ser-162 adjacent to Asp-161 completely abolished YegI kinase activity further validating the importance of DSD loop in YegI autophosphorylation.

Two possible mechanisms of kinase activation have been described: direct activation by ligand-promoted dimerization or indirect activation. The first evidence of ligand-promoted dimerization and activation of the kinase came from the structure of *M. tuberculosis* PknB (15,38) which forms back-to-back dimers of the N-terminal lobe of the catalytic domain that facilitates autophosphorylation. This mechanism of ligand-induced dimerization and activation was also observed for two other eSTKs, *M. tuberculosis* PknD (39), and *Pseudomonas aeruginosa* PpkA (40). In cases where the kinase lacks a ligand binding domain, an indirect mechanism has been proposed as a model for activation. For example, the structure of the PknB kinase domain alone with an ATP competitive inhibitor revealed formation of asymmetric front-to-front dimers via interaction between the αG helix and an ordered activation loop on one domain and αG helix and a disordered activation loop of the second domain. Substitutions in the αG helix interface resulted in decreased autophosphorylation suggesting that αG helix in the C-terminal lobe of the kinase is important for activation. The conformation of the two proteins in the dimer is such that one monomer acts like an activator while the other monomer acts like a substrate very similar to a transphosphorylation model (28). This provides an alternate mechanism of activation especially for soluble kinases that lack a membrane bound domain. Unlike most eSTKs, YegI lacks an extra cytoplasmic sensor domain and instead is a double pass membrane protein with both the N and C terminus in the cytoplasm (Fig. 6B). Additionally, our studies show that YegI follows second order kinetics which is consistent with dimer formation (Fig. 4A,B). Future studies will be aimed at understanding the mechanism of activation of YegI.

An interesting feature of the genomic organization of *yegI* is that it differs in closely related *E. coli* strains (Supplementary Fig.1). In most pathogenic strains and lab B strains of *E. coli, yegI* occurs immediately downstream of *pphC*, a PP2C-like phosphatase with a four nucleotide overlap while in K strains this locus is disrupted by a gene, *yegJ*, encoding an ORF without similarity previously identified proteins, suggesting potential regulatory differences between strains. In REL606 strains, *yegI* expression was detected only in stationary phase but at very low levels (data not shown). However, since the identity of the physiological stimulus that initiates *yegI* expression and activation remains unknown, it is difficult to interpret the functional implications of the altered gene locus.

An additional open question is the identity of YegI substrate(s). There are many candidates as ~100 cellular proteins were found to be phosphorylated on Ser/Thr residues in *E. coli* (4). A useful strategy to identify authentic in vivo substrates is comparative phosphoproteomics where the phosphoproteomes of a strain lacking either the eSTK or partner eSTP are compared with the WT strain. This methodology has been utilized previously for identifying phospho substrates of *B. subtilis* PrkC using deletion strains of eSTK PrkC and eSTP PrpC (41). This methodology has also been successfully utilized in *M. pneumoniae*, where 4 out of the 63 phosphorylated proteins were specifically enriched in a deletion strain of novel eSTK PrkC (42). Therefore, phosphoproteomic analysis might serve as a useful tool in identifying phosphorylated substrates of YegI by examining △*yegI* strains.

## Materials and Methods

### Bacterial strains and growth conditions

*E. coli* DH5α cells were used for regular cloning and *E. coli* LOBSTR (BL21-DE3) strains were used for expression of recombinant proteins. *E. coli* cells for expression were grown in LB Lennox broth supplemented with ampicillin (100 μg/ml) at 37 °C with shaking (220 rpm) unless otherwise specified. Genomic DNA from *E. coli* REL606 was isolated using a Wizard genomic DNA purification kit (Promega) following manufacturer’s instructions. Details of strains, plasmids and primers used in the study are described in Supplementary Tables S1, S2, and S3, respectively.

### Cloning and expression of YegI and YegI-NTD

The *yegI* gene was PCR amplified using Phusion polymerase (Thermo Scientific) from *E. coli* REL606 genomic DNA using primers (KR58/ KR59). Sequence for an N-terminal His6 tag was included in the primer. The PCR product was digested with *NcoI/SphI* and ligated into similarly digested pBAD24 backbone. Ligation products were transformed in DH5α cells and selected on LB/ampicillin plates. The resulting plasmid pKR31 generated an N-terminal His_6_-tagged YegI fusion protein. Plasmid cloning was subsequently verified by restriction enzyme digest and DNA sequencing (Operon).

For protein expression, plasmid was transformed into *E. coli* LOBSTR (Kerafast) cells and plated on LB ampicillin (100 μg/ml) agar plates. Single colonies were inoculated into 3 ml LB supplemented with ampicillin (100 μg/ml) for overnight cultures. The next day, a dilution of 1:250 in 500 ml LB was grown to an OD_600_ of 0.6-0.8. YegI recombinant protein was induced by addition of arabinose (0.2% w/v) for 3 h at 18 °C. Cells were harvested at 6000 x g for 15 min at 4 °C. Pellets were washed with ice-cold 50 mM EDTA and centrifuged at 7000 rpm for 15 min at 4 °C. Washed pellets were saved at −80 °C until use.

### Oligonucleotide site directed mutagenesis for YegI point mutants

Point mutation of lysine residue K39 was generated by two step overlap PCR mutagenesis using primer pairs (KR58/KR246) and (KR245/59) with Phusion polymerase. A second PCR was performed with primers (KR58/KR59) using primary PCR products as a template and the subsequent PCR products were digested with *Nco1/Sph1* and ligated into similarly digested pBAD24 to generate an N-terminal His_6_ tagged K39D YegI fusion protein. Plasmid cloning was subsequently verified by restriction enzyme digest and DNA sequencing (Operon).

Point mutation of aspartic acid residue D141 was generated by two step overlap PCR mutagenesis using primer pairs (KR58/ KR248) and (KR247/KR59) with Phusion polymerase. A second PCR was performed with primers (KR58/KR59) using primary PCR products as a template and the subsequent PCR products were digested with *Nco1/Sph1* and ligated into similarly digested pBAD24 to generate an N-terminal His_6_ tagged D141N YegI fusion protein. Plasmid cloning was subsequently verified by restriction enzyme digest and DNA sequencing (Operon).

Phosphoablative point mutants used in the study were generated using the same strategy as above and details of plasmid construction are described in Supplementary table S2.

### Purification of recombinant YegI

Frozen pellets were suspended in lysis buffer (50 mM Tris pH 8.0, 200 mM NaCl, 10 mM β-mercaptoethanol, 1 mM PMSF and 2% w/v sarkosyl) and incubated at room temperature for overnight lysis. Cells were subsequently lysed using sonication. Lysates were cleared by centrifugation at 15000 x g for 30 min at 4 °C and incubated in Pierce 5 ml columns with Ni-NTA agarose beads (Qiagen) at 4 °C for 1 h. Lysate was allowed to flow through and beads were washed 10 column volumes of wash buffer (50 mM Tris pH 8.0, 200 mM NaCl, 30 mM imidazole, 10 mM β-mercaptoethanol, 0.05% w/v sarkosyl). His tagged protein was eluted using 300 mM imidazole in 50 mM Tris pH 8.0, 200 mM NaCl, 10 mM β-mercaptoethanol and 0.05% w/v sarkosyl. Elution fractions were tested on 12% SDS-PAGE gel and fractions containing protein were pooled and dialyzed overnight at 4 °C using Snakeskin dialysis tubing 10K MWCO (Thermo Scientific) in kinase storage buffer (20 mM Tris pH 8.0, 125 mM NaCl, 10% glycerol, 1 mM DTT). Dialyzed protein was assessed for purity using 12% SDS-PAGE gel and stored at −80 °C. YegI variant proteins were purified using the same protocol. Proteins were stored at −80 °C.

### Autophosphorylation assays

#### Autophosphorylation assay using YegI point mutants

Autophosphorylation of YegI and point mutants was performed by addition of 5 μCi of [*γ*-^32^P] ATP (Perkin Elmer) to 0.2 μM of purified YegI or YegI point mutants in 10 μl of reaction buffer containing 50 mM Tris pH 7.5, 50 mM KCl, 0.5 mM DTT, 200 μM ATP, 10mM MgCl_2_ and 10mM MnCl_2_. Reactions were incubated at 37 °C for 30 mins and were stopped using 3X Laemmli buffer and boiled for 5 min at 95 °C. Samples were resolved on a 12% SDS-PAGE gel and visualized by staining with Coomassie dye. Radioactive gels were dried for 30 mins at 80 °C in a gel dryer, exposed to a phosphoscreen and visualized by autoradiography using a Typhoon Scanner (GE Healthcare).

#### Inhibition of autophosphorylation by staurosporine

Staurosporine sensitivity experiments were performed using 1 μM of YegI in 10 μl of reaction buffer containing 50 mM Tris pH 7.5, 50 mM KCl, 0.5 mM DTT, 10 mM MgCl_2_ and 10 mM MnCl_2_. Staurosporine (Calbiochem) was dissolved in DMSO to a obtain a stock concentration of 10 mM. Staurosporine stock (10 mM) was diluted in DMSO and added to the above kinase reaction to obtain final concentrations in the range of 12.5-100 μM. DMSO was added to the no inhibitor control. Reactions were incubated at 25 °C for 10mins followed by addition of 5 μCi of [*γ*-^32^P] ATP(Perkin Elmer). Reactions were incubated for additional 30 mins at 37 °C and were stopped using 3X Laemmli buffer. Samples were boiled for 5 min at 95 °C and resolved on a 12% SDS-PAGE gel. Gel was visualized by staining with Coomassie dye. Radioactive dried gel was exposed and visualized by autoradiography using a Typhoon Scanner (GE Healthcare).

#### Autophosphorylation assay using different bivalent cations

Autophosphorylation of YegI was performed by addition of 5 μCi of [*γ*-^32^P] ATP (Perkin Elmer) to 1 μM of purified YegI in 10 μl of reaction buffer containing 50 mM Tris pH 7.5, 50 mM KCl, 0.5 mM DTT, 200 μM ATP. MgCl_2_ or MnCl_2_ was added to a final concentration of 0.01, 0.1,1 and 10 mM. Reactions were incubated at 37 °C for 30 mins and were stopped using 3X Laemmli buffer and boiled for 5 min at 95 °C. Samples were resolved on a 12% SDS-PAGE gel and visualized by staining with Coomassie dye. Radioactive gels were dried for 30 mins at 80 °C in a gel dryer, exposed to a phosphoscreen and visualized by autoradiography using a Typhoon Scanner (GE Healthcare). To examine autophosphorylation of YegI mutants (S160FD161G and D161G), reactions were carried out in reaction buffer containing either 10 mM MgCl_2_ or 10 mM MnCl_2_.

#### Autophosphorylation assay to examine kinetics of YegI

Autophosphorylation of YegI was performed by addition of 5 μCi of [*γ*-^32^P] ATP (Perkin Elmer) to different concentrations of purified YegI (0.0625,0.125,0.25,0.5,1,2 μM) in 10 μl of reaction buffer containing 50 mM Tris pH 7.5, 50 mM KCl, 0.5 mM DTT, 200 μM ATP, 10mM MgCl_2_ and 10mM MnCl_2_. Reactions were incubated at 37 °C for 30 mins and were stopped using 3X Laemmli buffer and boiled for 5 min at 95 °C. Samples were resolved on a 12% SDS-PAGE gel and visualized by staining with Coomassie dye. Radioactive gels were dried for 30 mins at 80 °C in a gel dryer, exposed to a phosphoscreen and visualized by autoradiography using a Typhoon Scanner (GE Healthcare).

### Membrane topology analysis

#### Construction of YegI PhoA-lacZα fusion and derivatives

To study the membrane topology of YegI, plasmid pKR72 was constructed that encodes a dual *pho-lac* reporter in-frame to full length YegI. The YegI-PhoA-lacZα was constructed by a PCR-mediated overlap extension (44). First, the *yegI* gene fragment encoding aa 2-646 was PCR amplified using Phusion polymerase (Thermo Scientific) from *E. coli* REL606 genomic DNA using primers (KR223/KR224). Alkaline phosphatase PhoA fragment (aa 21-471) and the alpha fragment of beta galactosidase LacZ (aa 5-60) was PCR amplified from *E. coli* MG1655 genomic DNA using primers (KR225/KR226) for *phoA* and (KR227/KR228) for *lacZα* respectively. Eighteen nucleotides encoding a GSGSGS linker along with eighteen nucleotides from the *phoA* DNA was introduced in the reverse primer KR224 to generate an overlapping region between *yegI* and *phoA*. Similarly, eighteen nucleotides from the *lacZ* DNA was introduced in the reverse primer KR226 to generate an overlapping region between *phoA* and *lacZ*. The *yegI* and *phoA* PCR products were purified and used as templates for a second PCR to create a fused *yegI-linker-pho* DNA that was amplified using primers (KR229/KR226). Sequence for N-terminal His6 tag was included in the primer KR229. Purified PCR products *yegI-linker-phoA* DNA and *lacZ* DNA were used as templates for third and final round of PCR to create a fused *yegI-linker-pho-lac* fusion that was amplified using primers (KR229/KR228). The PCR product was digested with *Nhe1/Kpn1* and ligated into similarly digested pBAD24 plasmid backbone. Ligation products were transformed into DH5α cells and selected on LB/ampicillin plates. The resulting plasmid pKR72 generated an N-terminal His6-tagged YegI-PhoA-lacZα fusion protein. Plasmid cloning was subsequently verified by restriction enzyme digest and DNA sequencing (Operon).

Plasmid derivatives of pKR72 with pho-lac fusions were constructed in frame with different codons of YegI using the Q5 Site-Directed mutagenesis kit (NEB). Primers used to generate pho-lac fusions in frame to different codons E439 (KR230/KR231), T432 (KR230/KR232), R414 (KR230/KR233), E390 (KR230/KR234), E340 (KR230/KR235) were designed using NEBaseChanger. Primers were designed such that the 5’ ends were annealing back-to-back following manufacturer’s instructions. PCR amplification was performed using Q5 hotstart high fidelity master mix (NEB) with respective primers and 2 ng of Plasmid pKR72 as template DNA. The PCR product was subsequently treated with Kinase/Ligase/Dpn1 mix for 5 mins and transformed in NEB5alpha competent *E. coli* cells (NEB C2987). This method generated plasmids with in-frame *pho-lac* fusions to different codons of YegI at positions E439 (pKR73), T432 (pKR74), R414 (pKR75), E390 (pKR76), E340 (pKR77). pKR78, a derivative of pKR72 lacking yegI was used as a control plasmid and was constructed using the same method using primers KR230/KR240. Plasmid cloning was subsequently verified by colony PCR and DNA sequencing (Operon).

#### Membrane topology assay

To assess membrane topology *in vivo*, *E. coli* DH5α cells were transformed with the resulting pho-lac fusion plasmids and streaked on dual indicator plates containing LB agar with 5-bromo-4-chloro-3-indolyl phosphate disodium salt (X-Phos) (Sigma) at a final concentration of 80μg/ml and 6-chloro-3-indolyl-β-D-galactoside (Red-Gal) (Sigma) at a final concentration of 100 μg/ml as indicators. L-arabinose at a final concentration of 0.1% was added as an inducer for reporter expression. 100μg/ml of ampicillin was added for plasmid selection. Plates were incubated at 25° C for 24 h and scanned using EPSON image scanner.

## Acknowledgements

We would like to thank present and former members of our lab for helpful discussions. We would like to thank Elizabeth Nagle for performing initial experiments characterizing YegI. This work was supported by an HHMI International Student Fellowship to KR and by NIH grant GM114213 to JD.

## Author contributions

KR performed all of the experiments and KR and JD wrote the manuscript.

## Supplementary figure legends

**Figure S1:**
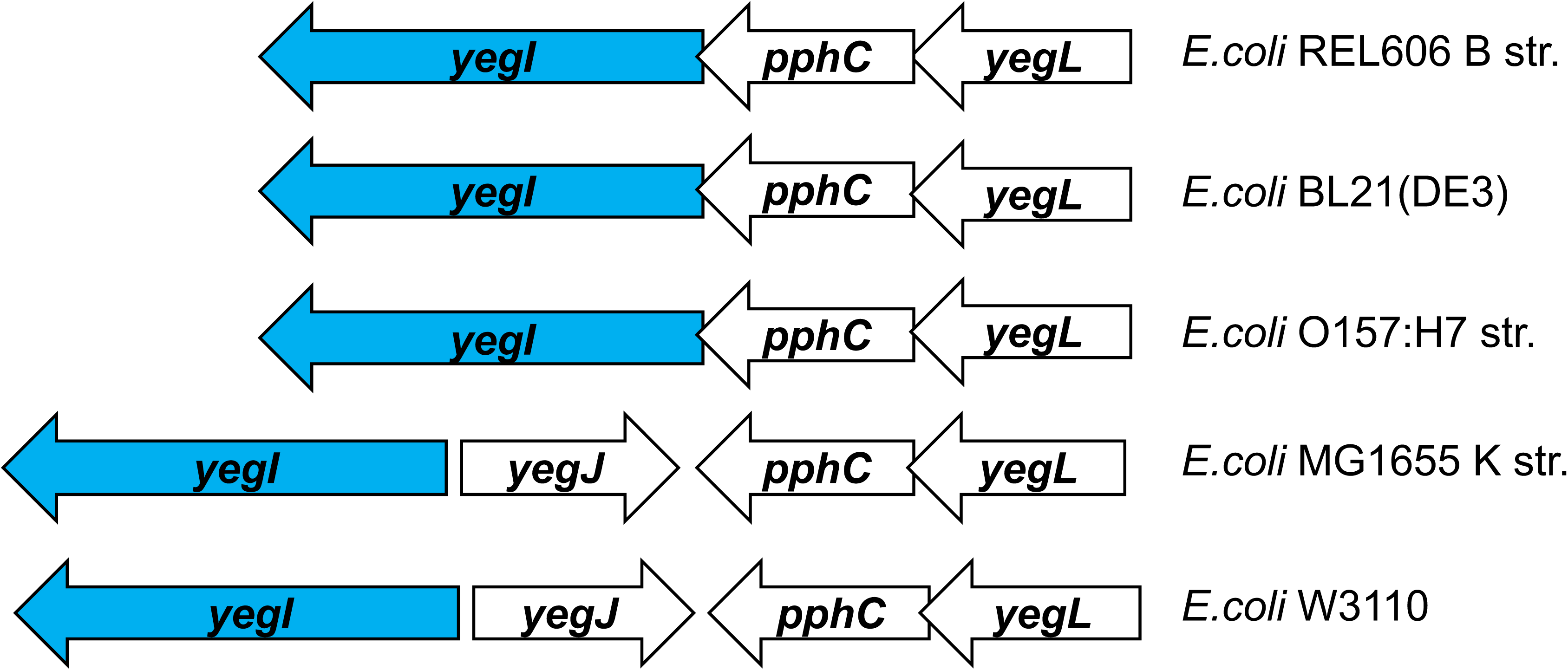
Genome organization of *yeg* operon in different *E. coli* strain backgrounds. Thick arrows denote orfs. Genes *yegL, pphC, yegI* encode a predicted Von Willebrand factor, a PP2C-like phosphatase and an eukaryotic-like Ser/Thr kinase respectively. The *yegJ* gene encodes a protein of unknown function. *E. coli* strain backgrounds are indicated on the right.

**Figure S2:**
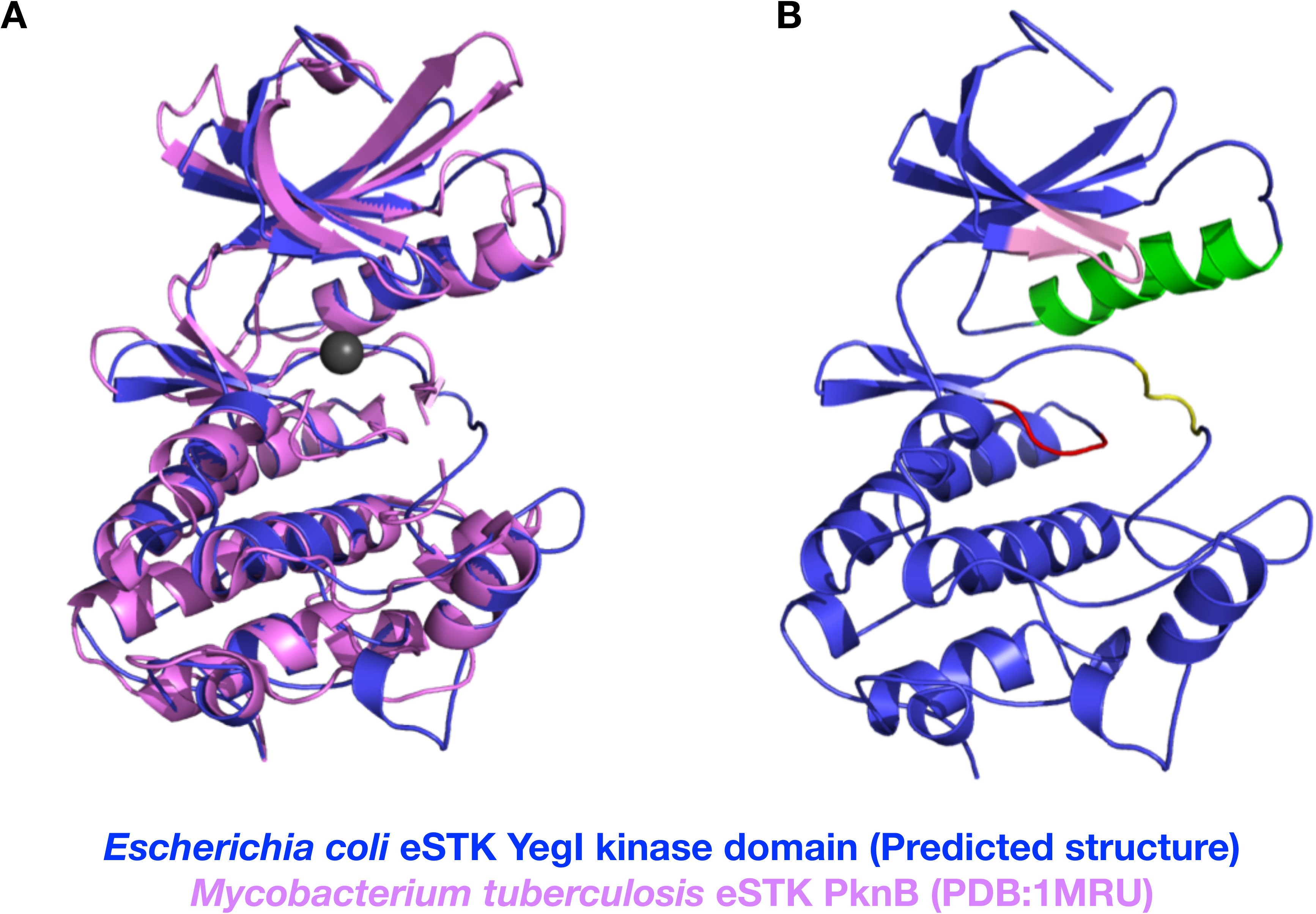
Predicted structure of YegI kinase domain (aa 1-302). **(A)** Structural overlay of predicted structure of YegI kinase domain (blue) and *Mycobacterium tuberculosis* eSTK PknB (pink) (Protein data bank PDB:1MRU). Mg^2+^ ion is represented as a grey sphere. **(B)** SWISS MODEL algorithm (1) was used to predict structural model of YegI kinase domain. Structurally important motifs such as P-loop (pink), Helix C (green), Catalytic loop (red), activation loop (yellow) and P+1 loop (black) are indicated in structure. Corresponding amino acid sequences are depicted in Fig 1.

**Figure S3:**
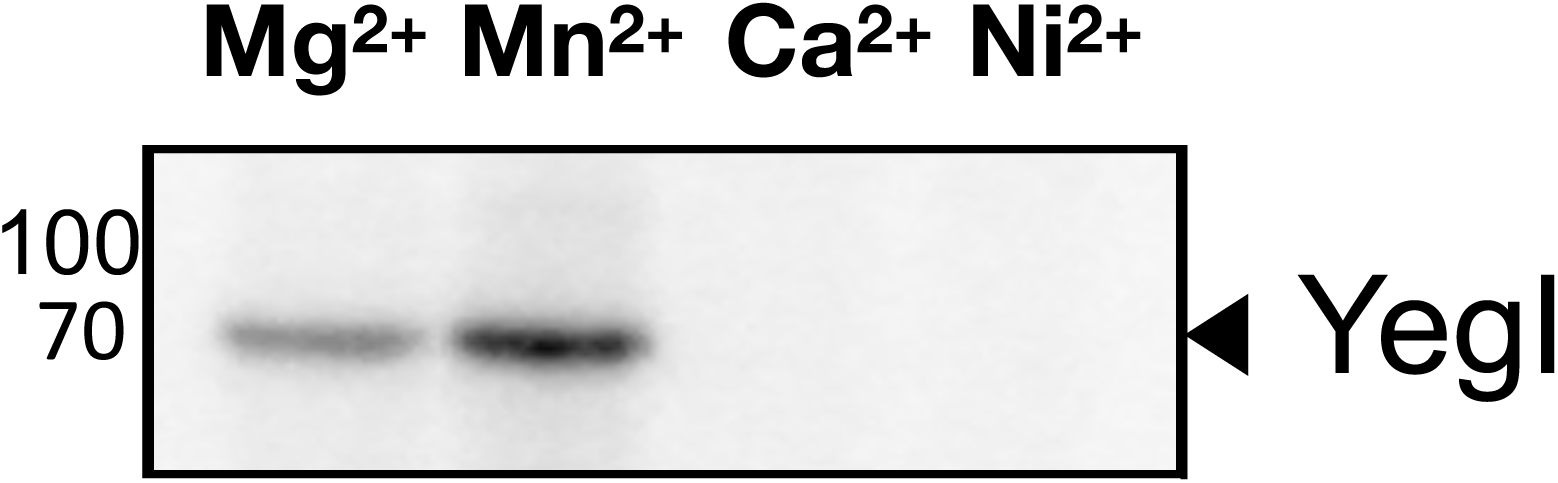
Requirement of bivalent cations. Autophosphorylation reactions were carried out at 37 °C with 1 µM of YegI (WT) in kinase buffer with either 10 mM of MgCl_2_/ MnCl_2_, CaCl_2_/ NiCl_2_. Reactions were stopped at t=30 mins and run on 12% SDS-PAGE followed by autoradiography

**Table S1:**
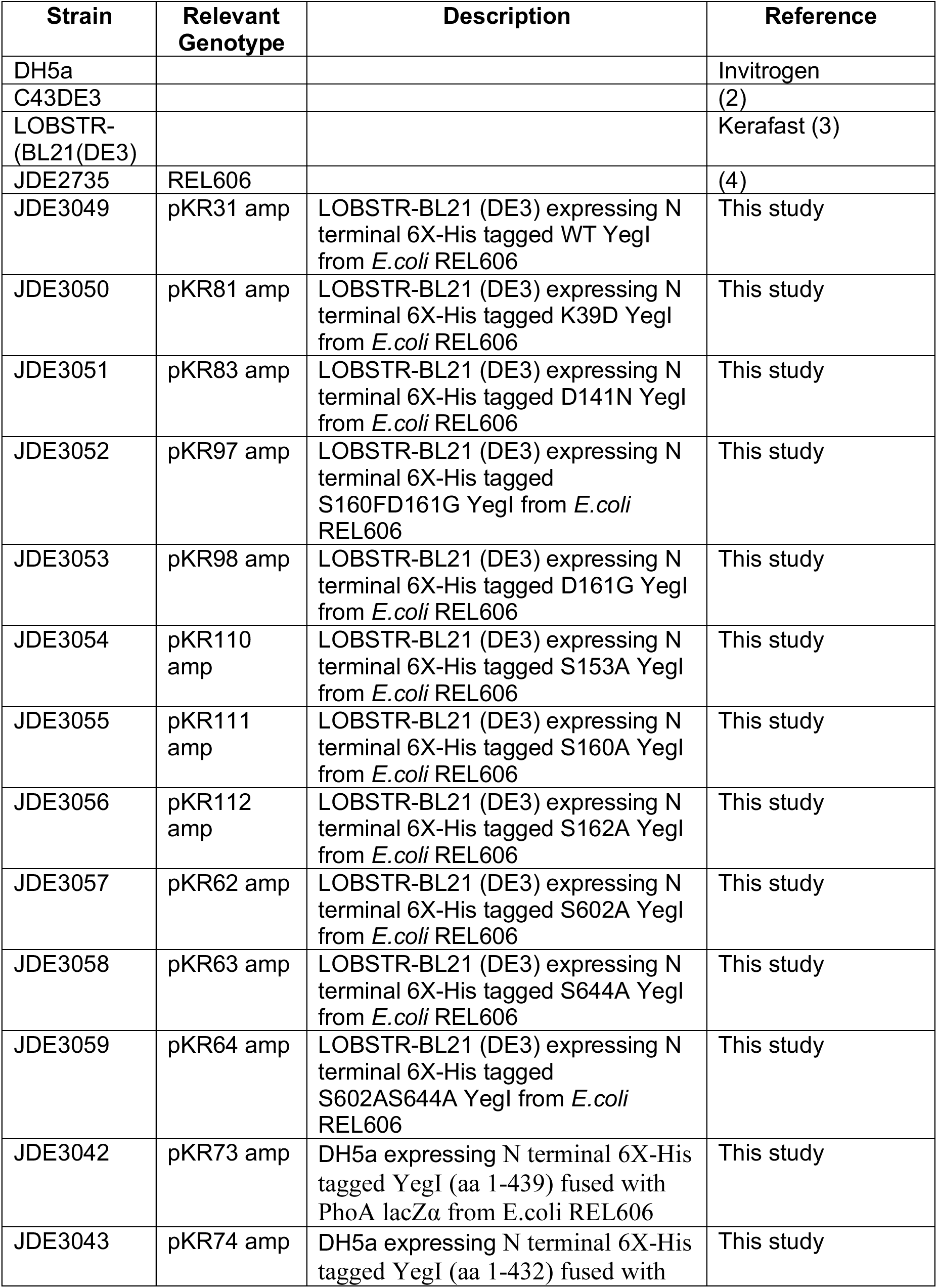
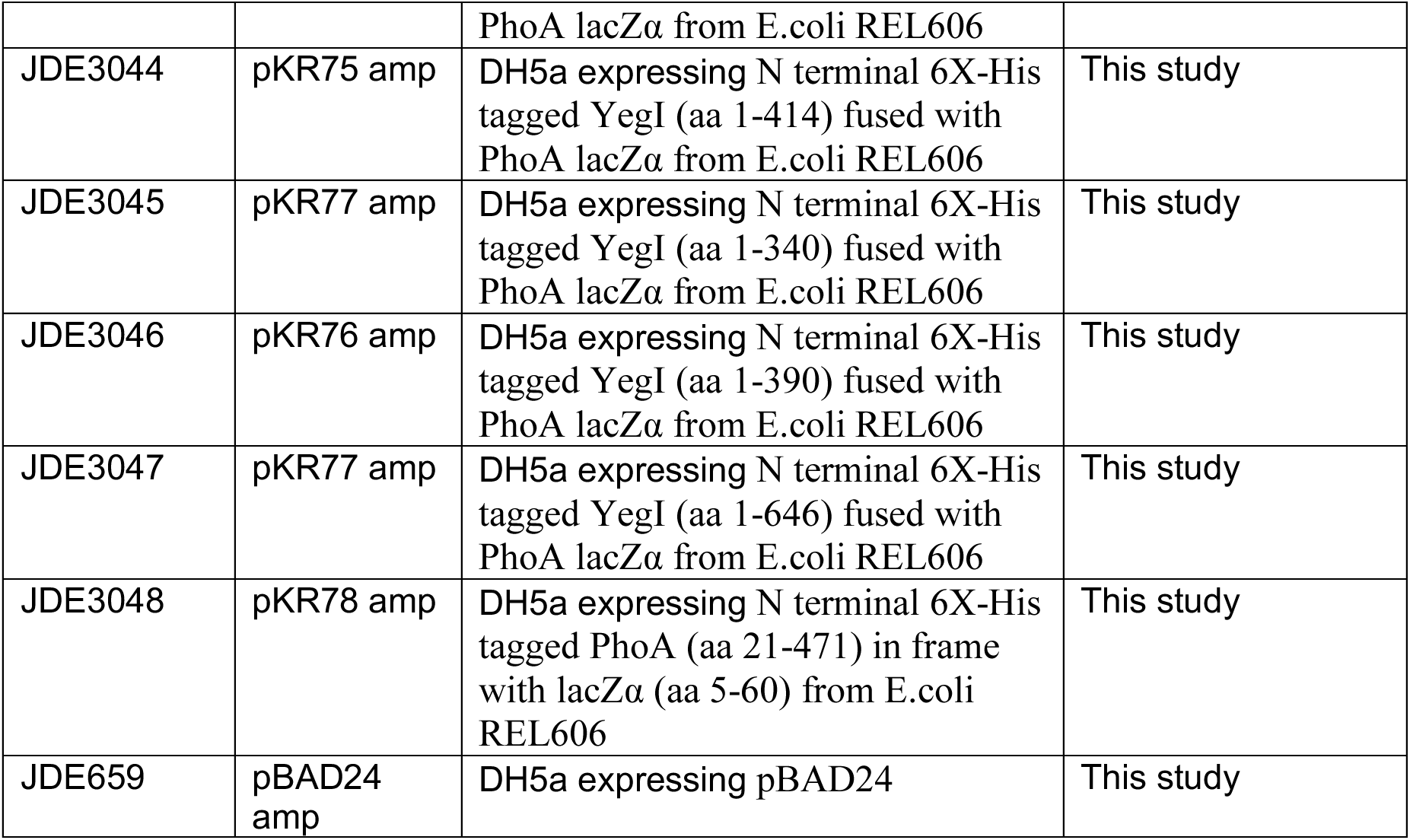
Bacterial Strains

**Table S2:** Plasmids

| **Plasmid Name** | **Description** | **Reference** |
| --- | --- | --- |
| pBAD24 | Cloning vector, ampR | (5) |
| pKR31 | yegl was amplified from REL606 genomic DNA using KR58/KR59 primers, digested with NcoI/SphI and ligated into pBAD24 | This study |
| pKR81 | REL606 K39D yegl was generated by two step PCR using KR58/KR246 and KR245/KR59 primer pairs followed by amplification with KR58/KR59. Resulting product was digested with NcoI/SphI and ligated into pBAD24 | This study |
| pKR83 | REL606 D141N yegl was generated by two step PCR using KR58/KR248 and KR247/KR59 primer pairs followed by amplification with KR58/KR59. Resulting product was digested with NcoI/SphI and ligated into pBAD24 | This study |
| pKR97 | REL606 S160FD161G yegl was generated by Q5 site directed mutagenesis using KR285/KR286 and pKR31 as a template. | This study |
| pKR98 | REL606 D161G yegl was generated by Q5 site directed mutagenesis using KR285/KR286 and pKR31 as a template. | This study |
| pKR110 | REL606 S153A yegl was generated by two step PCR using KR58/KR158 and KR157/KR59 primer pairs followed by amplification with KR58/KR59. Resulting product was digested with NcoI/SphI and ligated into pBAD24 | This study |
| pKR111 | REL606 S160A yegl was generated by two step PCR using KR58/KR306 and KR305/KR59 primer pairs followed by amplification with KR58/KR59. Resulting product was digested with NcoI/SphI and ligated into pBAD24 | This study |
| pKR112 | REL606 S162A yegl was generated by two step PCR using KR58/KR308 and KR307/KR59 primer pairs followed by amplification with KR58/KR59. Resulting product was digested with NcoI/SphI and ligated into pBAD24 | This study |
| pKR62 | REL606 S602A yegl was generated by two step PCR using KR58/KR153 and KR152/KR59 primer pairs followed by amplification with KR58/KR59. Resulting product was digested with NcoI/SphI and ligated into pBAD24 | This study |
| pKR63 | REL606 S644A yegl was generated by two | This study |
|  | step PCR using KR58/KR155 and KR154/KR59 primer pairs followed by amplification with KR58/KR59. Resulting product was digested with NcoI/SphI and ligated into pBAD24 |  |
| pKR64 | REL606 S602AS644A yegI was generated by two step PCR using pKR62 as template and KR58/KR155 and KR154/59 primer pairs followed by amplification with KR58/KR59. Resulting product was digested with NcoI/SphI and ligated into pBAD24 | This study |
| pKR72 | REL606 *yegI-phoA-lacZ* fusion was generated by overlap extension PCR using KR223/KR224 , KR225/KR226, KR227/KR228 primer pairs followed by amplification with KR229/KR228. Resulting product was digested with NheI/KpnI and ligated into pBAD24 | This study |
| pKR73 | REL606 YegI E439-phoA lacZ was generated by Q5 site directed mutagenesis using KR230/KR231 and pKR72 as a template. | This study |
| pKR74 | REL606 YegI T432-phoA lacZ was generated by Q5 site directed mutagenesis using KR230/KR232 and pKR72 as a template. | This study |
| pKR75 | REL606 YegI R414-phoA lacZ was generated by Q5 site directed mutagenesis using KR230/KR233 and pKR72 as a template. | This study |
| pKR76 | REL606 YegI E390-phoA lacZ was generated by Q5 site directed mutagenesis using KR230/KR234 and pKR72 as a template. | This study |
| pKR77 | REL606 YegI E340-phoA lacZ was generated by Q5 site directed mutagenesis using KR230/KR235 and pKR72 as a template. | This study |
| pKR78 | Control phoA lacZ was generated by Q5 site directed mutagenesis using KR230/KR240 and pKR72 as a template. | This study |

**Table S3:**
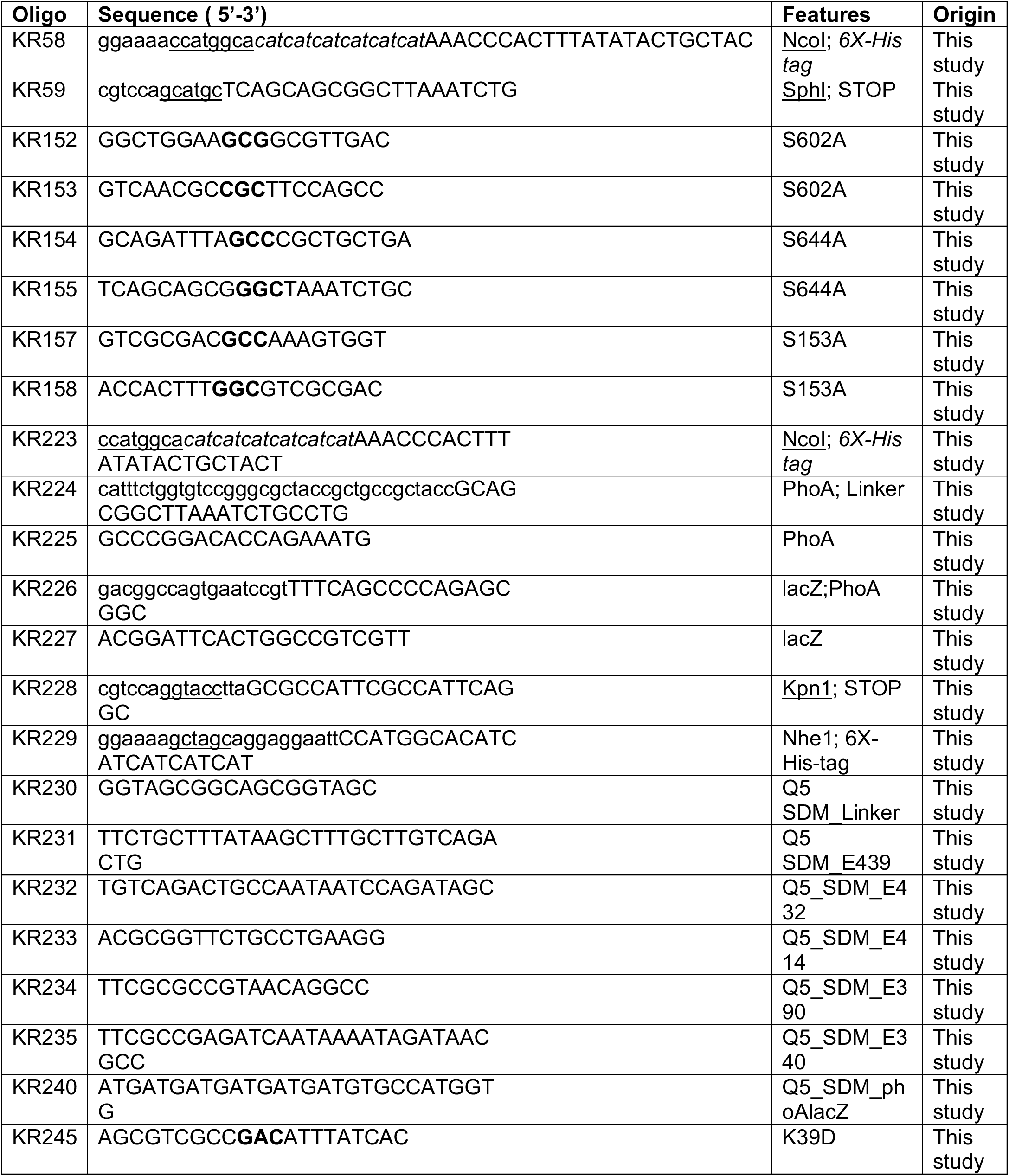
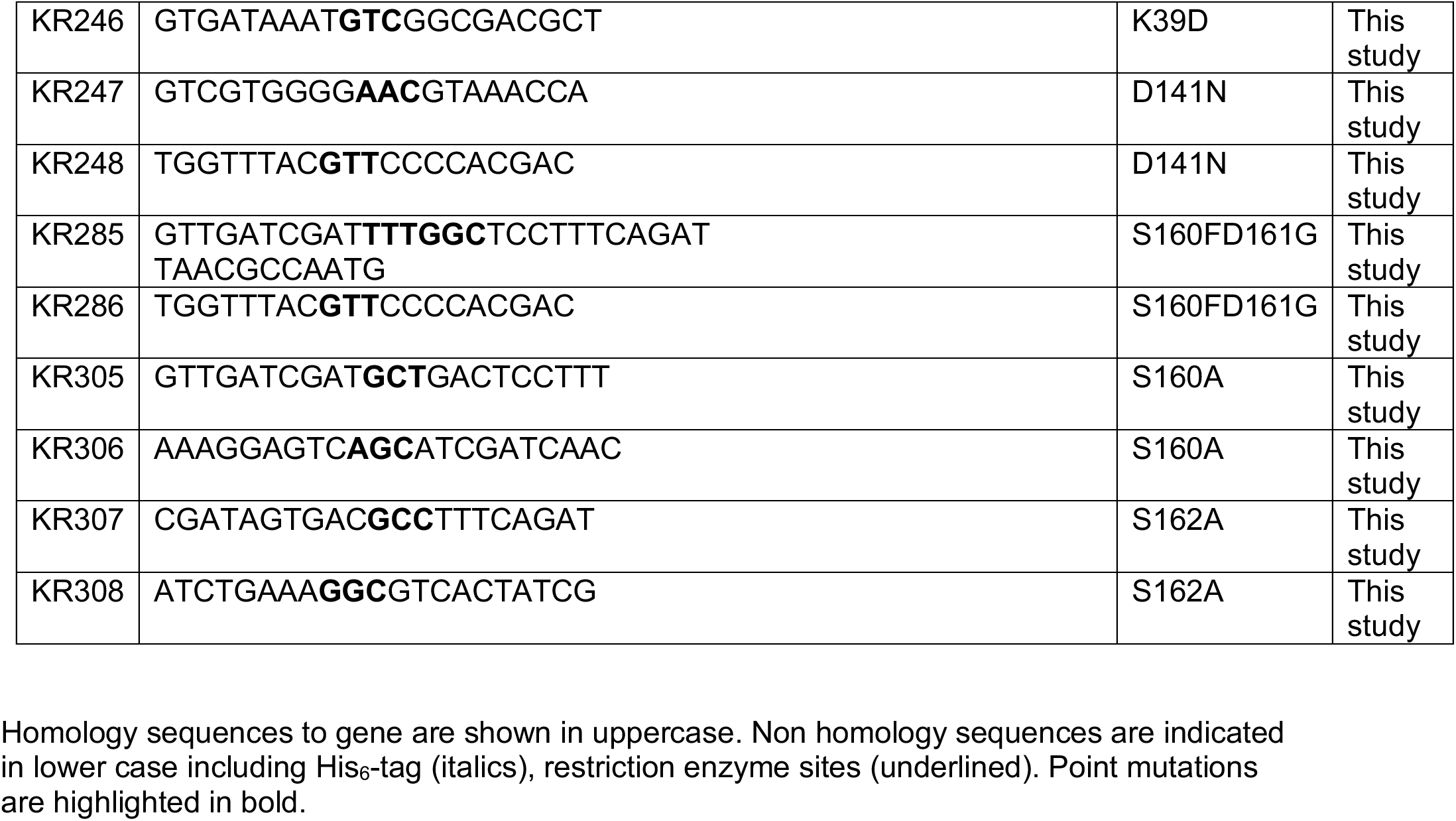
Primers

